# A near chromosome-scale genome assembly of the Common pine sawfly (*Diprion pini*, Linnaeus, 1758)

**DOI:** 10.64898/2026.03.19.712881

**Authors:** Saskia Wutke, Craig Michell, Carita Lindstedt

**Author notes:** Correspondence: Saskia Wutke, Ecology and Genetics research unit, University of Oulu, Oulu, Finland.

## Abstract

The common pine sawfly, *Diprion pini*, is a widespread defoliator of pine forests across Europe and Asia, with outbreaks causing substantial ecological and economic damages. However, genomic resources for this species have been limited, hindering advances in molecular ecology or pest management. Here, we present a near chromosome-level reference genome for *D.pini*, generated using PacBio HiFi reads, Oxford Nanopore MionION long reads, and 10x Genomics linked reads. The final assembly is organized into mostly chromosome-sized scaffolds. It spans a length of 268 Mb, comprises 81 scaffolds, and has a scaffold N50 of 18.7 Mb. BUSCO analysis (hymenoptera_odb10) indicates a high genome completeness of 97.2%. With 22,7 kb the mitochondrial genome is unusually large due to an extended non-coding control region (6,874 bp). Gene prediction identified 26,335 protein-coding genes, of which 12,769 were functionally annotated. Comparative analyses with other sawflies and Apocrita identified 2,472 proteins unique to *D. pini*, some of which are putatively associated with the processing of plant secondary metabolites. Notably, our genome assembly highlights that, when a closely related, high-quality reference genome is available, chromosome-scale assemblies can be generated without the need of Hi-C sequencing. The genome provides a valuable foundation for the development of improved monitoring and management strategies for *D. pini* outbreaks and contributes to advancing fundamental research on Hymenoptera evolution.

## Introduction

The common pine sawfly, *Diprion pini* (Hymenoptera: Diprionidae), is a widespread defoliator of coniferous forests across Europe and Asia, with outbreaks causing extensive damage to commercially and ecologically important pine forests (Barre et al., 2002; De Somviele et al., 2004). However, despite its ecological importance, genomic resources for this species remain scarce, hindering advances in molecular ecology or pest management. As a member of the paraphyletic Symphyta (sawflies and woodwasps), *D. pini* occupies a phylogenetic position that is critical for understanding the early evolution of Hymenoptera, the insect order that includes both the mainly phytophagous sawflies and the typically parasitoid or eusocial Apocrita (ants, bees, and wasps). In general, sawflies represent an understudied lineage within Hymenoptera, which limits comparative genomics across this order such as identifying ancestral genomic features (He et al., 2025; Zhang et al., 2025) or reconstructing the evolution of parasitism (Polaszek and Vilhemsen, 2023) and sociality (Ritter et al., 2026). Recent efforts to manage *D. pini* outbreaks have focused on mechanical methods like handpicking the larvae, as well as the use of chemical insecticides (Sevim and Sevim, 2024). These methods are labor-intensive and increasingly constrained by environmental concerns. New genomic tools, facilitated by advances in genome sequencing technologies, can support and inform pest management. Yet the lack of a high-quality reference genome for *D. pini* limits progress in identifying genetic targets for pest control, genomic monitoring of outbreak source and spread, or exploring adaptations to climate-driven stressors.

Here, we present the first near chromosome-level reference genome assembly and structural annotation for *Diprion pini*, generated using PacBio long-read sequencing, supplemented with and ONT MionION long reads, and 10x linked-read scaffolding. Additionally, we explore syntenic relationships and gene homology with other Diprionidae genomes. This resource provides a foundation for advancing genomic-driven pest management strategies such as the development of molecular assays for early detection and monitoring, and genome-informed strategies for mitigating outbreak impacts. Beyond its benefit for applied forest entomology, the *D. pini* genome will be useful for broader comparisons between Symphyta and Apocrita at the level of genome architecture or gene content. Pine sawflies are also becoming a new model system to study herbivore-host plant co-evolution (Bagley et al., 2017; Linnen and Farrell, 2010), protective coloration (Koskenpato et al., 2026; Lindstedt et al., 2022; Linnen et al., 2018) and cooperative behavior (Lindstedt et al., 2025, 2018; Ritter et al., 2026).

## Materials and Methods

### DNA extraction and sequencing

*Diprion pini* imagines were reared from cocoons collected from soil searches in Northwest Poland in autumn 2021. High molecular weight (HMW) genomic DNA (gDNA) was isolated following the Chromium™ Genome Demonstrated Protocol for HMW gDNA extraction from single insects. For each extraction, the whole insect was ground with a pestle and digested overnight at 37℃ in lysis buffer (333 mM NaCl, 100 mM EDTA pH 8, 10 mM Tris HCl pH 8, 0,5% SDS, and 100 µg/ml Proteinase K). After adding 5M NaCl, the mixture was centrifuged, and the aqueous phase transferred to a new tube. HMW gDNA was precipitated in ice-cold 100% ethanol and collected by centrifugation. The pellet was air-dried and dissolved thoroughly in 0.1× TE buffer. The eluted DNA was quantified using a Qubit 3.0 Fluorometer (Thermo Fisher Scientific, USA). Quality was assessed via agarose gel electrophoresis (1.0%) and NanoDrop spectrophotometric analysis (Thermo Fisher Scientific), with A260/A280 ratios between 1.8–2.0 and A260/A230 > 2.0.

For preparation of sequencing libraries, the DNA of two individuals was used. PacBio HiFi long-read libraries were prepared from HMW gDNA of a female and sequenced on the PacBio Sequel II platform at the DNA Sequencing and Genomics Laboratory (BIDGEN) of the University of Helsinki. DNA extracted from a male specimen was used for preparing a 10x Chromium linked-read library (10x Genomics, USA) and a long-read library for ONT MinION (Oxford Nanopore Technologies, UK). The linked-read library was prepared using the Chromium Genome Reagent Kits v2 (10x Genomics, USA) and sequenced on the Illumina NovaSeq platform at the Institute for Molecular Medicine Finland (FIMM) of the University of Helsinki after size-selecting the DNA for fragments over 10 kb on a BluePippin (Sage Science, USA). For ONT MinION library preparation we used the Rapid Barcoding Sequencing (SQK-RBK004, Oxford Nanopore Technologies, UK) according to manufacturer’s instructions. The prepared library was sequenced on a R9.4.1 MinION flow cell (FLO-MIN106, Oxford Nanopore Technologies, UK).

### Genome survey, assembly, and quality assessment

Genome size and repetitive content were estimated based on a k-mer approach, in which the k-mer (k = 21) distribution count was generated by jellyfish v2.3.1 (Marçais and Kingsford, 2011) using the 10x reads. Subsequently, the resulting histogram was analyzed with the GenomeScope2 web interface (Vurture et al. 2017, http://genomescope.org/genomescope2.0/) with no maximum k-mer coverage (max k-mer coverage = −1). The genome was assembled with Hifiasm v0.19 (Cheng et al., 2022, 2021) using default parameters and PacBio HiFi and ONT MinION long reads as input. We used purge_dups v1.2.6 (Guan et al., 2020) to remove duplicated regions caused by contig overlaps and haplotigs. Potential misassemblies in the purged genome were resolved using RagTag v2.1.0 “correct” (Alonge et al., 2022) with the genome of *Diprion similis* (GCF_021155765.1) as a reference due to its taxonomic proximity (Wutke et al., 2024). Subsequently, the corrected contigs were placed into scaffolds during three scaffolding steps: 1) with scaff10x (https://github.com/wtsi-hpag/Scaff10X) using the 10x linked reads, 2) with LINKS v2.0.1 (Warren et al., 2015) using both PacBio HiFi and MinION long reads, and 3) with RagTag v2.1.0 “scaffold” (Alonge et al., 2022) which ordered, oriented, and grouped scaffolds into pseudo-chromosomes based on synteny with the *D. similis* reference.

The quality of each stage of the assembly process was assessed with BUSCO v5.4.7 (Manni et al., 2021b, 2021a; Simão et al., 2015) using the hymenoptera_odb10 dataset, and by calculating k-mer completeness and QV estimates using Meryl dbs (k = 19) and Merqury v1.3 (Rhie et al., 2020). The assembly was further analyzed with the BlobToolKit suite (Challis et al., 2020) to ensure the absence of contamination.

The initial mitochondrial genome was assembled with MitoHiFi v3.2.1 (Uliano-Silva et al., 2023) using the PacBio HiFi data. This assembly was used for extracting mitochondrial reads by mapping the HiFi reads back to the assembly with minimap2 (Li, 2021) and re-assembling the mapped reads using Hifiasm v0.19 (Cheng et al., 2022, 2021). We also mapped the short reads to the new assembly to check mapping depth for the control region as a recent study concluded that long mitochondrial genomes may be more common than previously thought (Olli et al., 2024). The annotation of the mitogenome was done with the MITOS2 (Bernt et al., 2013) web application on the Galaxy Europe server (https://usegalaxy.eu/). Some of the gene annotations done by MITOS2 were corrected based on alignments with mitogenome sequences of related diprionid species (from GenBank).

### Structural and functional genome annotation

Repeat annotation was conducted with the Extensive de-novo TE Annotator (EDTA) v2.2.0 (Ou et al., 2019) which combines commonly used TE identification programs such as LTRharvest (Ellinghaus et al., 2008), LTR_FINDER (Xu and Wang, 2007), LTR_retriever (Ou and Jiang, 2018), HelitronScanner (Xiong et al., 2014), and RepeatModeler (A. F. A. Smit et al., 2015) to compile a comprehensive de-novo TE library. The final repeat library was used for soft-masking the genome with RepeatMasker v4.1.2-p1 (A. F. A. Smit et al., 2015). The RepeatMasker script calcDivergenceFromAlign.pl was used to collect the Kimura distances for each TE class for generating a repeat landscape plot. We also used tidk (Brown et al., 2025) to identify telomers by exploring the presence of telomeric repeats within the assembled pseudo-chromosomes and unplaced scaffolds. We searched the scaffolds for the repeat motif AACCT which is the canonical form of the common arthropod repeat (TTAGG)_n_ (Brown et al., 2025; Stoianova et al., 2025).

Gene prediction was performed on the soft-masked genome with the BRAKER2 annotation pipeline (Brůna et al., 2021), using both mapped RNA and protein evidence. The RNA-seq data originates from the transcriptome-based phylogeny of Hymenoptera by Peters et al. (2017, <u>SRR1503030</u>). RNA reads were mapped to the genome with STAR v2.7.11a (Dobin et al., 2013) and the resulting bam file was used as input for BRAKER. The protein evidence is based on the arthropod-specific dataset of the OrthoDB v11 (Kuznetsov et al., 2023). The prediction results from both runs were combined with TSEBRA v1.1.2.5 (Gabriel et al., 2021) with default settings.

For functional annotation we first inferred orthologous groups in *D. pini*, five other diprionid species (*D. similis* GCF_021155765.1, *Neodiprion fabricii* GCF_021155785.1, *N. lecontei* GCF_021901455.1, *N. pinetum* GCF_021155775.1, *N. virginiana* GCF_021901495.1), two other sawflies (*Orussus abietus* GCF_000612105.2, *Athalia rosae* GCF_917208135.1) and three Apocrita (*Atta cephalotes* GCF_000143395.1, *Apis mellifera* GCF_003254395.2, *Polistes dominula* GCF_001465965.1) with OrthoFinder v2.5.5 (Emms and Kelly, 2019, 2015). The complete set of proteins for *D. pini* predicted by BRAKER as well as the unique set of orthologs identified by OrthoFinder were then subjected to annotation in the PANNZER2 web server (Törönen and Holm 2022, http://ekhidna2.biocenter.helsinki.fi/sanspanz/), a fully automated tool that predicts Gene Ontology (GO) annotations and functional descriptions (DE). The 30 most common GO terms in each set were visualized as word clouds.

### Comparative genomics

Synteny analysis was conducted by comparing hymenoptera_odb10 single-copy orthologs identified by BUSCO v5.4.7 (Manni et al., 2021b, 2021a; Simão et al., 2015) across the focal genome and two related species (*D.similis*, *Neodiprion lecontei*). Conserved syntenic blocks were identified following the protocol for visualizing syntenies between genomes (Manni, Berkeley, Seppey, Simão, et al. 2021, https://gitlab.com/ezlab/busco_protocol) and plotted with the R package RIdeogram (Hao et al., 2020).

Visualization of shared orthogroups (identified by OrthoFinder) was performed with UpSetR v1.4.0 (Conway et al., 2017; Lex et al., 2014) with complex set intersections quantified through the upset() function.

Whole-genome alignment against the *D. similis* genome was performed using nucmer (MUMmer4 package, Marçais et al. 2018). The alignment was processed with show-coords to generate collinear regions. A dot plot of the alignment was created with the R script “mummerCoordsDotPlotly.R” (https://github.com/tpoorten/dotPlotly).

## Results and Discussion

### Genome assembly and quality assessment

We generated a highly contiguous, near chromosome-level assembly of *Diprion pini* using PacBio HiFi (47× coverage) and ONT MinION (2×) long reads, scaffolded with 10x linked reads. The final assembly (= Hifiasm’s primary haplotype assembly) spans 268 Mb (contig N50: 7 Mb; scaffold N50: 18.7 Mb, Fig. 1, Table 1, Supplementary information), consistent with genome sizes reported for other symphytan lineages, with 97.2% of the hymenoptera_odb10 BUSCO genes (5,991 genes) recovered as complete (Fig. 2A). The GenomeScope analysis predicted a haploid genome size of 253 Mb and a repeat content of 19.6% (Fig. 2B). Although both are close to the final estimates, the difference between the predicted and the final values may stem from the use of long HiFi reads in the assembly, which likely improve the resolution of repetitive sequences. Merqury analysis confirmed high base accuracy (QV 61.3 i.e., ca. 1 error in each Mb) and k-mer completeness (94.2%), reflecting a highly accurate consensus sequence with few unrepresented genomic regions. This assembly is among the most contiguous and complete sawfly genomes to date, attributable to Hifiasm’s haplotype-resolved assembly and hybrid scaffolding.

**Fig. 1.**
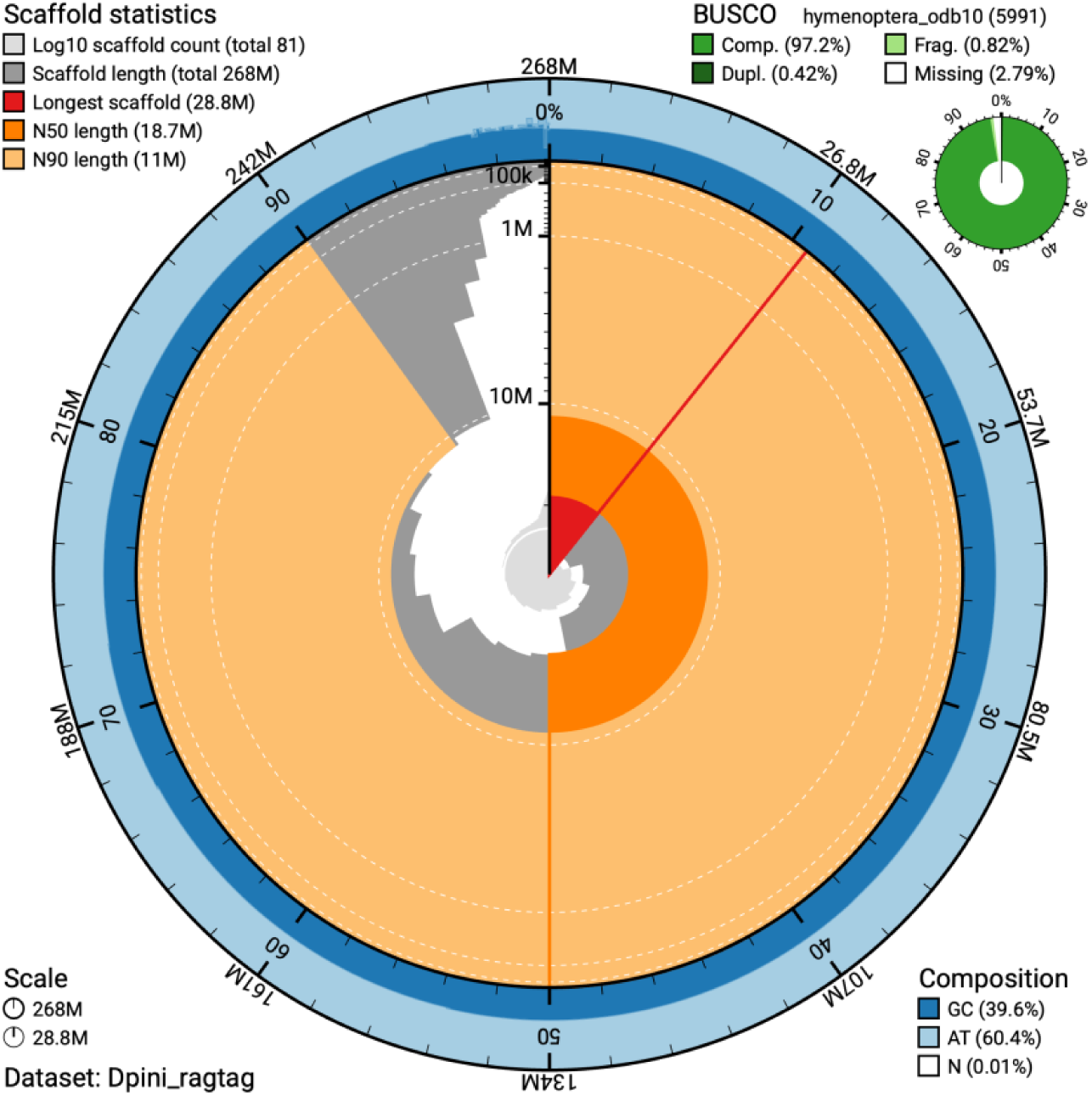
Snail plot visualization of the assembly statistics created with BlobToolKit with scaffold statistics in the top left corner. The red line marks the longest scaffold.

**Fig. 2.**
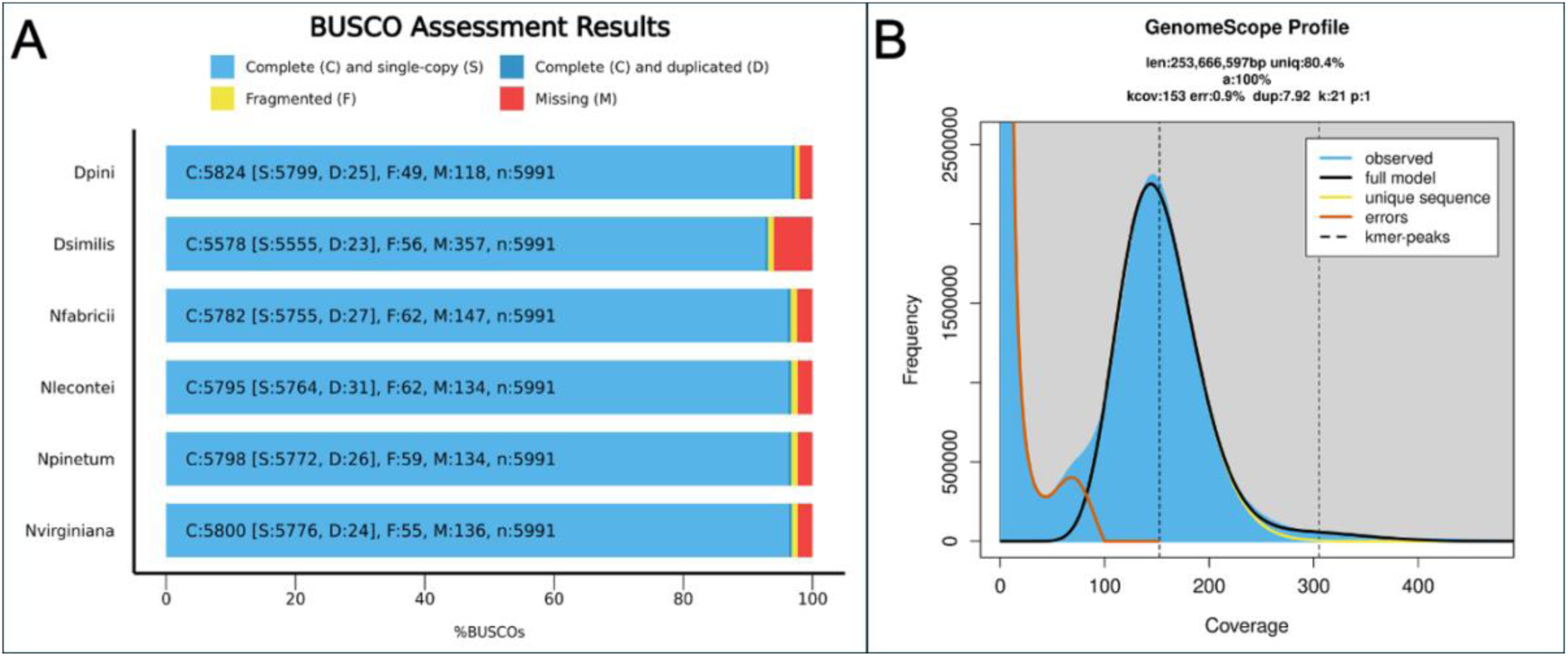
Genome survey and quality metrics for the *D. pini* reference genome. (A) BUSCO completeness based on the hymenoptera_odb10 database for the newly generated *D. pini* genome compared to published genomes of other Diprionidae species. (B) GenomeScope profile of 21-mer analysis using paired-end reads as input.

**Table 1.**
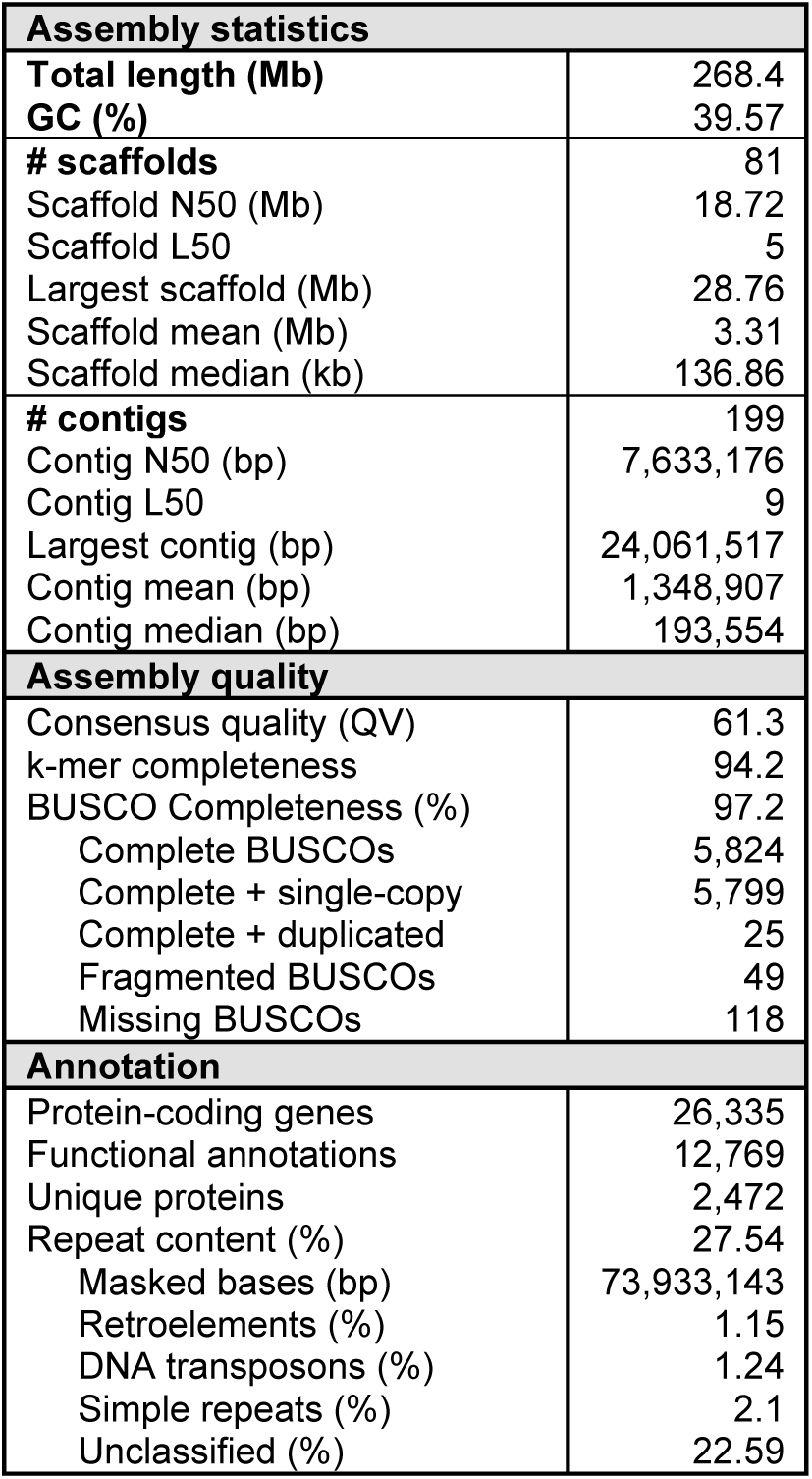
Assembly and annotations statistics.

| **Assembly statistics** |  |
| --- | --- |
| **Total length (Mb)** | 268.4 |
| **GC (%)** | 39.57 |
| **# scaffolds** | 81 |
| Scaffold N50 (Mb) | 18.72 |
| Scaffold L50 | 5 |
| Largest scaffold (Mb) | 28.76 |
| Scaffold mean (Mb) | 3.31 |
| Scaffold median (kb) | 136.86 |
| **# contigs** | 199 |
| Contig N50 (bp) | 7,633,176 |
| Contig L50 | 9 |
| Largest contig (bp) | 24,061,517 |
| Contig mean (bp) | 1,348,907 |
| Contig median (bp) | 193,554 |
| **Assembly quality** |  |
| Consensus quality (QV) | 61.3 |
| k-mer completeness | 94.2 |
| BUSCO Completeness (%) | 97.2 |
| Complete BUSCOs | 5,824 |
| Complete + single-copy | 5,799 |
| Complete + duplicated | 25 |
| Fragmented BUSCOs | 49 |
| Missing BUSCOs | 118 |
| **Annotation** |  |
| Protein-coding genes | 26,335 |
| Functional annotations | 12,769 |
| Unique proteins | 2,472 |
| Repeat content (%) | 27.54 |
| Masked bases (bp) | 73,933,143 |
| Retroelements (%) | 1.15 |
| DNA transposons (%) | 1.24 |
| Simple repeats (%) | 2.1 |
| Unclassified (%) | 22.59 |

The mitochondrial genome (22.7 kb, Fig. 3) was fully resolved, exhibiting a conserved gene order typical of Bileteria (Lavrov 2007). With almost 23 kb the mitochondrial genome is unusually large compared to most other animal mitogenomes, which can be attributed to an unusually long non-coding control region (6,874 bp). Read-depth inspection supported the authenticity of this region, rather than it being an assembly artifact. Yet, large mitochondrial genomes might be more the norm than the exception. As Olli et al. (2024) suggested, the control region length might be underestimated for many organisms due to mapping and assembly issues using short Illumina reads that do not spam the whole control region. With steadily improving long-read sequencing technologies, such large mitogenome sizes are being reported more and more frequently (Olli et al., 2024).

**Fig. 3.**
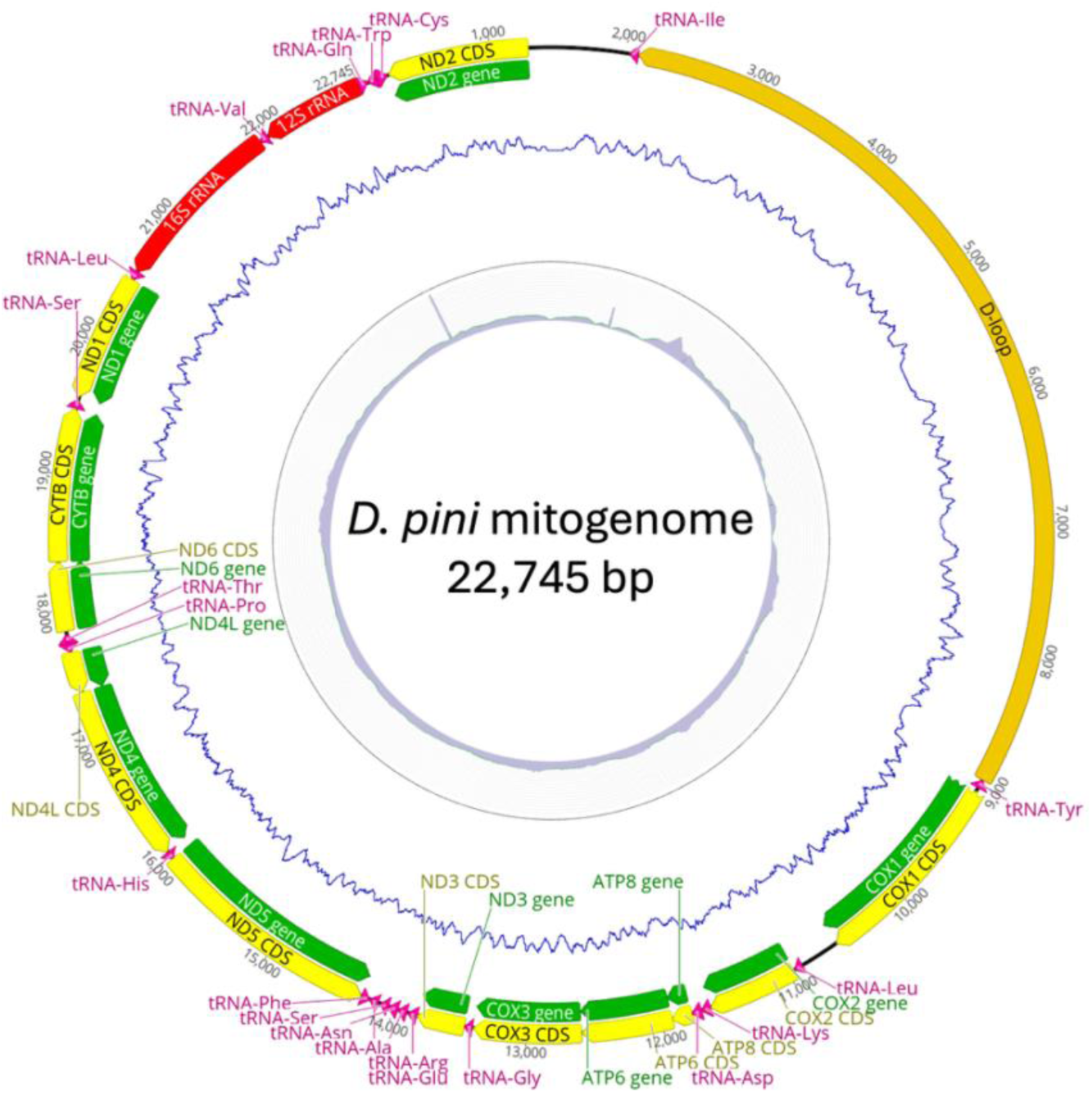
Mitochondrial genome map of *Diprion pini*. Inner histogram represents coverage (up to 13,000×) of linked reads and blue line shows GC content.

### Structural and functional genome annotation

Repetitive elements accounted for 27.54% of the genome (Table 1, Supplemantary Fig. S4), of which most were unclassified repeats (22.59%). Comparative data from hymenopteran genomes indicate that repeat abundance is variable across the order (Sproul et al., 2023). While the genomes of several representatives of the basally branching Symphyta show comparatively large repeat fractions relative to many apocritan genomes (Sproul et al., 2023), the pattern is heterogeneous. However, comparing repeat content across genome assemblies generated with different technologies and pipelines is delicate, because much of the apparent variation might be technical rather than biological. Therefore, comprehensive taxon sampling and consistent repeat annotation pipelines are required to determine whether elevated repeat content is a general feature of ancestral hymenopteran genomes or the result of lineage-specific TE dynamics. The tidk analysis showed that four of the 14 assembled pseudo-chromosomes were enriched in the telomeric repeat motif on both scaffold ends, and another nine were enriched on one end of the scaffold. Due to their repetitiveness, assembling telomeres can be challenging. Nevertheless, the presence of the repeat motif near several scaffold ends supports the near-complete chromosome-level scaffolding of our assembly (Supplementary Fig. S5).

Gene prediction with BRAKER2 produced a comprehensive gene set of 26,335 protein-coding genes that further suggests a high quality of our assembly. Of these predicted genes, 12,769 could be functionally annotated with PANNZER2. The comparison of orthologs with other Hymenoptera identified 7,073 shared orthogroups, likely forming the conserved core set of proteins of the lineage. A further 648 orthogroups containing 2,472 proteins were unique to *D. pini* (Fig. 4). While the functions of many proteins remain unknown or relate to general processes, several carry annotations suggesting roles in interaction with host plants or natural enemies (e.g., toxin activity, GO:0090729, or modulation of process of another organism, GO: 0035821, Fig. 5, Supplemantary Fig. S6). These annotations may indicate involvement in managing plant defenses e.g., suppressing the production of plant secondary metabolites or sequestering these compounds for the insect’s own protection against predators. Such proteins represent promising targets for investigating the genetic basis of adaptive traits, including the digestion of terpene-rich pine foliage and physiological responses to environmental stress.

**Fig. 4.**
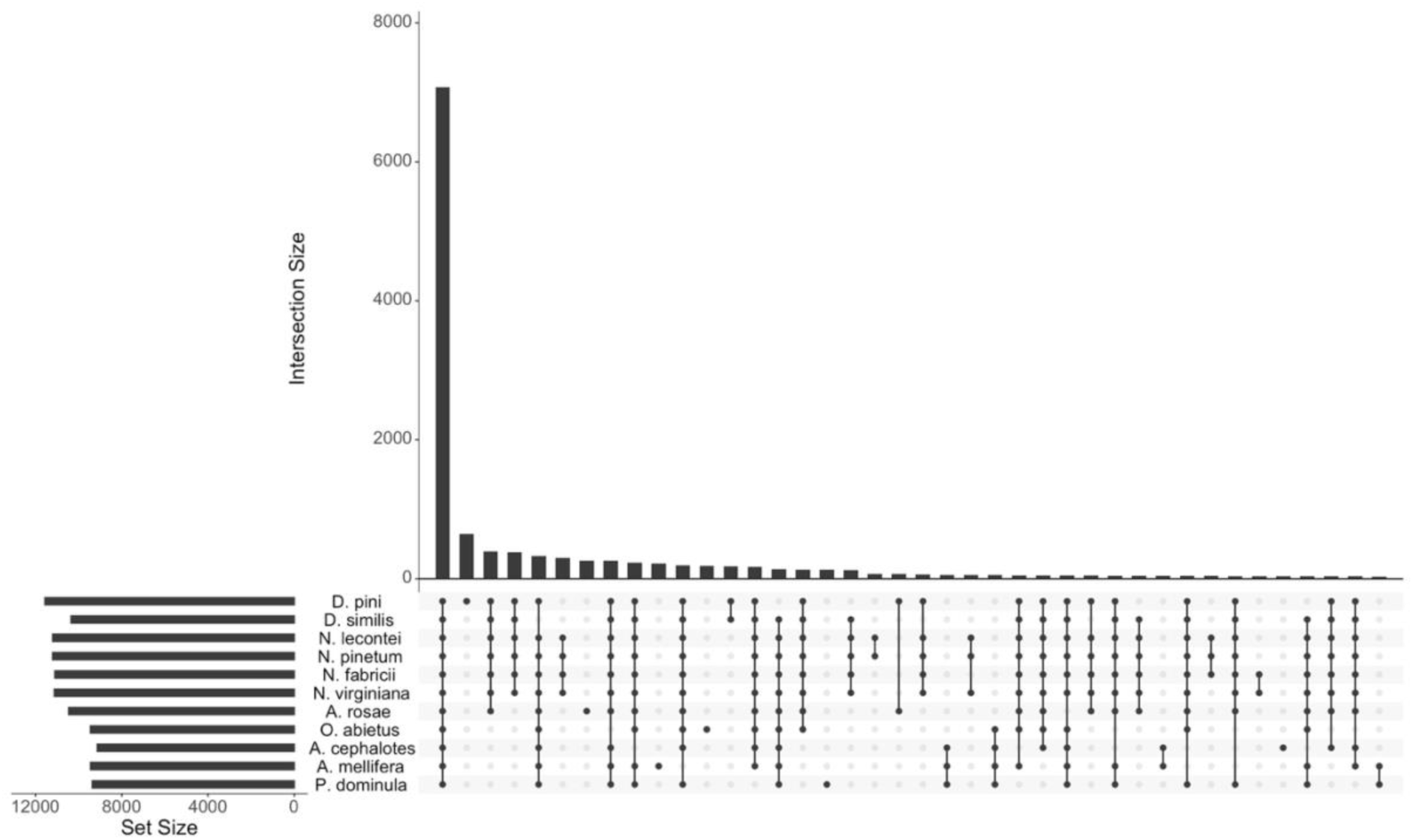
UpSet plot showing the intersection of orthogroups identified by OrthoFinder across the 11 hymenopteran protein sets included here. Horizontal bars (= set sizes) indicate the total number of orthogroups found in each species’ genome, while vertical bars represent the number of orthogroups shared by a given intersection of species. Dots in the lower panel indicate the species in each intersection with single dots representing orthogroups unique to a particular species, and dots connected by lines representing orthogroups shared by several species.

**Fig. 5.**
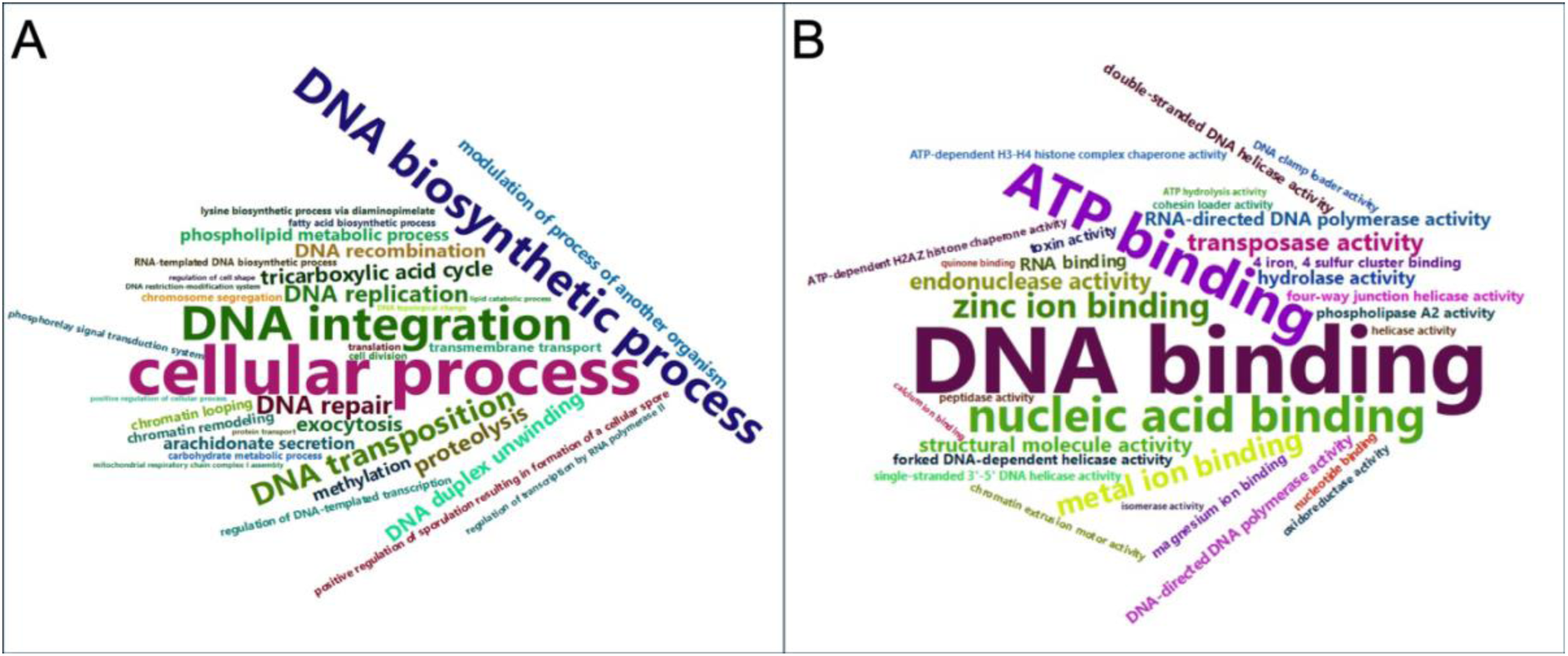
Word clouds of the 30 most abundant gene ontology (GO) terms in the *D. pini* unique protein set. (A) Annotations of biological processes (BP); (B) Annotations of molecular functions (MF).

### Comparative genomics

Whole-genome alignments revealed a high degree of synteny between *Diprion pini* and *Neodiprion lecontei* (Fig. 6, Supplemantary Fig. S7), a representative of another genus within Diprionidae. This strong conservation of syntenic blocks, despite the twofold difference in haploid chromosome number (n = 7 in *Neodiprion* vs. n = 14 in *Diprion*), suggests that large chromosomal segments have remained intact during karyotype evolution. Thus, the divergence in chromosome number likely reflects chromosome fusion or fission events (Robertsonian changes) rather than large-scale intrachromosomal rearrangement, a pattern reported across diverse insect lineages (De Vos et al., 2020; Huang et al., 2025; Nakatani and McLysaght, 2019). The observed synteny with *Diprion similis* (Fig. 6, Supplemantary Fig. S7) is not surprising given that the reference genome of that species was used to break misassemblies, order, and orient scaffolds of the *D. pini* assembly. *D. pini* and *D. similis* show the same chromosome number and almost identical morphology in their chromosomes (Rousselet et al., 1998). Based on that and the fact that the *D. similis* reference genome has been assembled to chromosome-level (Davis et al., 2023), it seems only logical to guide the scaffolding process with this assembly.

**Fig. 6.**
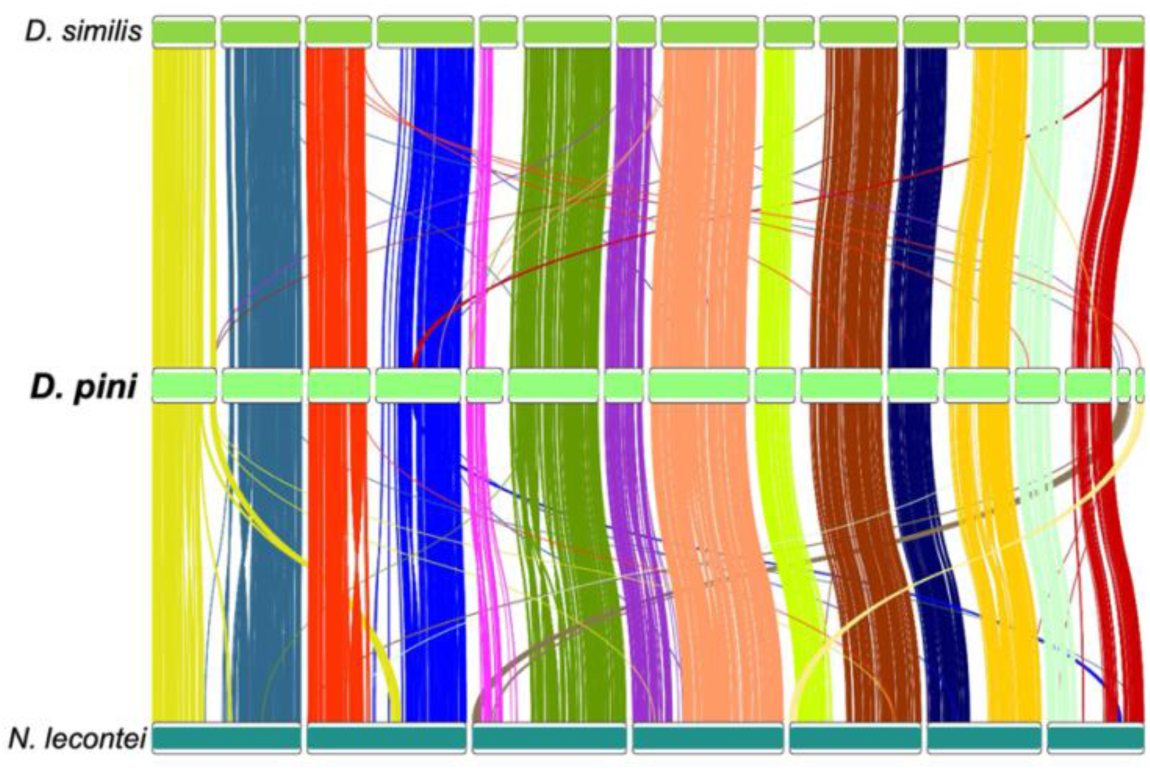
Synteny of BUSCO genes (hymenoptera_odb10) with *D. pini* in the middle, *D. similis* in the top and *N. lecontei* in the bottom.

The *D. pini* genome presented here represents a high-quality reference, expanding genomic resources for sawflies, a group historically under-represented in Hymenoptera genomics. Given the ecological and economic importance of *D. pini* as a major defoliator of pine forests, its genome constitutes a needed resource to develop more effective, targeted, and sustainable strategies for monitoring, predicting, and managing outbreaks of this important forest pest. At the same time, it offers opportunities for advancing fundamental entomological research e.g., evolutionary and behavioural ecology of sawflies. Importantly, our highly contiguous assembly achieved near-complete chromosome-level scaffolding. This demonstrates that the availability of a closely related, high-quality reference genome can enable the generation of chromosome-scale assemblies without relying on cost- and labor-intensive Hi-C sequencing. As genomic resources continue to grow across Diprionidae and other sawfly families, this assembly is a valuable contribution toward resolving longstanding questions concerning host specialization and diversification in early-diverging Hymenoptera.

## Supplementary Information

Table S1. Assembly statistics and quality metrics after each step of the assembly process.

Table S2. RepeatMasker output.

Figure S1. Workflow visualization of the genome assembly and annotation pipeline.

Figure S2. Workflow visualization for assembling and annotating the mitochondrial genome.

Figure S3. BlobToolKit assembly views. (A) Blob plot showing the distribution of assembly scaffolds on GC proportion and coverage; (B) Cumulative assembly span plot showing curves for subsets of scaffolds assigned to each phylum relative to the overall assembly.

Figure S4. Repeat landscape.

Figure S5. Occurrence of the common arthropod repeat (TTAGG)_n_ within the assembled scaffolds.

Figure S6. Word clouds of the 30 most abundant gene ontology (GO) terms in the *D. pini* complete protein set. (A) Annotations of biological processes (BP); (B) Annotations of molecular functions (MF).

Figure S7. Dot plots of whole-genome alignments: (A) *D. pini* vs. *D. similis*, (B) *D. pini* vs. *N. lecontei*.

## Data availability

All raw data and the assembly are available in the European Nucleotide Archive (ENA) under project PRJEB109745, with the following accessions: ERR16829677 (HiFi), ERRxxxxxxxx (MinION), ERR16829746 (10x), and GCA_982290575 (genome assembly). The annotation files are available from the Github repository via https://github.com/swutke/Diprion_pini_genome/.

## Code availability

All bioinformatic tools and software used for genome assembly, annotation, and data analysis in this study were operated strictly according to their official user manuals. Software versions and parameters are comprehensively documented in the Methods section or in the Supplementary information. All scripts and modified configuration files are also available from the Github repository https://github.com/swutke/Diprion_pini_genome/.

## Acknowledgements

We would like to thank Zbigniew Borowski for collecting and providing cocoon samples of *D. pini*.

## Authors’ contributions

C.L.: funding acquisition, writing—review and editing; C.M.: methodology, writing—review and editing; S.W.: conceptualization, data curation, formal analysis, funding acquisition, investigation, methodology, project administration, resources, validation, visualization, writing—original draft, writing—review and editing; all authors gave final approval for publication.

## Generative AI disclosure

The authors used ChatGPT-4 to assist with language editing. All content has been reviewed and revised by the authors.

