## Supplemental Tables and Figures for "A near chromosome-scale genome assembly of the Common pine sawfly (*Diprion pini*, Linnaeus, 1758)"

**Table S1.** Assembly statistics and quality metrics after each step of the assembly process.

| Step<br><br>Tool | Assembly (HiFi + ONT)<br><br>Hifiasm | Haplotig merging<br><br>purge_dups | Breaking misassemblies<br><br>RagTag | Scaffolding (linked)<br><br>Scaff10X | Scaffolding (HiFi + ONT)<br><br>LINKS | Order + orient scaffolds<br><br>RagTag |
|---|---|---|---|---|---|---|
| **Assembly statistics** | | | | | | |
| **Total length (Mb)** | 282.6 | 268.4 | 268.4 | 268.4 | 268.4 | 268.4 |
| **GC (%)** | 39.61 | 39.58 | 39.57 | 39.57 | 39.57 | 39.57 |
| **# scaffolds** | 203 | 82 | 267 | 221 | 150 | 81 |
| Scaffold N50 (Mb) | 9.04 | 9.42 | 7.63 | 8.7 | 8.7 | 18.72 |
| Largest scaffold (Mb) | 24.06 | 24.06 | 24.06 | 24.06 | 24.06 | 28.76 |
| Scaffold mean (Mb) | 1.39 | 3.27 | 1 | 1.21 | 1.79 | 3.31 |
| Scaffold median (kb) | 89.12 | 407.9 | 131.9 | 136.2 | 193.03 | 136.86 |
| **# contigs** | 203 | 82 | 267 | 267 | 199 | 199 |
| Contig N50 (bp) | 9,035,521 | 9,422,795 | 7,633,176 | 7,633,176 | 7,633,176 | 7,633,176 |
| Largest contig (bp) | 24,061,517 | 24,061,517 | 24,061,517 | 24,061,517 | 24,061,517 | 24,061,517 |
| Contig mean (bp) | 1,391,939 | 3,273,567 | 1,005,365 | 1,005,365 | 1,348,907 | 1,348,907 |
| Contig median (bp) | 89,119 | 407,934 | 131,895 | 131,895 | 193,554 | 193,554 |
| **Assembly quality** | | | | | | |
| Consensus quality (QV) | 60.7 | 61.3 | 61.3 | 61.3 | 61.3 | 61.3 |
| k-mer completeness | 95.3 | 94.2 | 94.2 | 94.2 | 94.2 | 94.2 |
| **BUSCO Completeness (%)** | 97.3 | 97.5 | 97.3 | 97.3 | 97.3 | 97.2 |
| Complete BUSCOs | 5,825 | 5,837 | 5,827 | 5,827 | 5,827 | 5,824 |
| Complete + single-copy | 5,636 | 5,803 | 5,784 | 5,784 | 5,785 | 5,799 |
| Complete + duplicated | 189 | 34 | 43 | 43 | 42 | 25 |
| Fragmented BUSCOs | 43 | 36 | 45 | 45 | 45 | 49 |
| Missing BUSCOs | 123 | 118 | 119 | 119 | 119 | 118 |

**Table S2.** RepeatMasker output.

bases masked: 73933143 bp ( 27.54 %)

| | number of elements* | length occupied | percentage of sequence |
|---|---|---|---|
| Retroelements | 3761 | 3087750 bp | 1.15 % |
| SINEs: | 0 | 0 bp | 0.00 % |
| Penelope: | 0 | 0 bp | 0.00 % |
| LINEs: | 1317 | 1144539 bp | 0.43 % |
| CRE/SLACS | 0 | 0 bp | 0.00 % |
| L2/CR1/Rex | 466 | 374197 bp | 0.14 % |
| R1/LOA/Jockey | 449 | 466714 bp | 0.17 % |
| R2/R4/NeSL | 0 | 0 bp | 0.00 % |
| RTE/Bov-B | 0 | 0 bp | 0.00 % |
| L1/CIN4 | 0 | 0 bp | 0.00 % |
| LTR elements: | 2444 | 1943211 bp | 0.72 % |
| BEL/Pao | 53 | 109489 bp | 0.04 % |
| Ty1/Copia | 639 | 224837 bp | 0.08 % |
| Gypsy/DIRS1 | 1745 | 1608266 bp | 0.60 % |
| Retroviral | 0 | 0 bp | 0.00 % |
| DNA transposons | 7068 | 3323797 bp | 1.24 % |
| hobo-Activator | 817 | 154234 bp | 0.06 % |
| Tc1-IS630-Pogo | 320 | 211157 bp | 0.08 % |
| En-Spm | 0 | 0 bp | 0.00 % |
| MULE-MuDR | 279 | 95547 bp | 0.04 % |
| PiggyBac | 0 | 0 bp | 0.00 % |
| Tourist/Harbinger | 4 | 419 bp | 0.00 % |
| Other (Mirage, P-element, Transib) | 0 | 0 bp | 0.00 % |
| Rolling-circles | 364 | 180362 bp | 0.07 % |
| Unclassified: | 181546 | 60638083 bp | 22.59 % |
| Total interspersed repeats: | | 67049630 bp | 24.98 % |
| Small RNA: | 0 | 0 bp | 0.00 % |
| Satellites: | 0 | 0 bp | 0.00 % |
| Simple repeats: | 144695 | 5624716 bp | 2.10 % |
| Low complexity: | 21791 | 1078435 bp | 0.40 % |

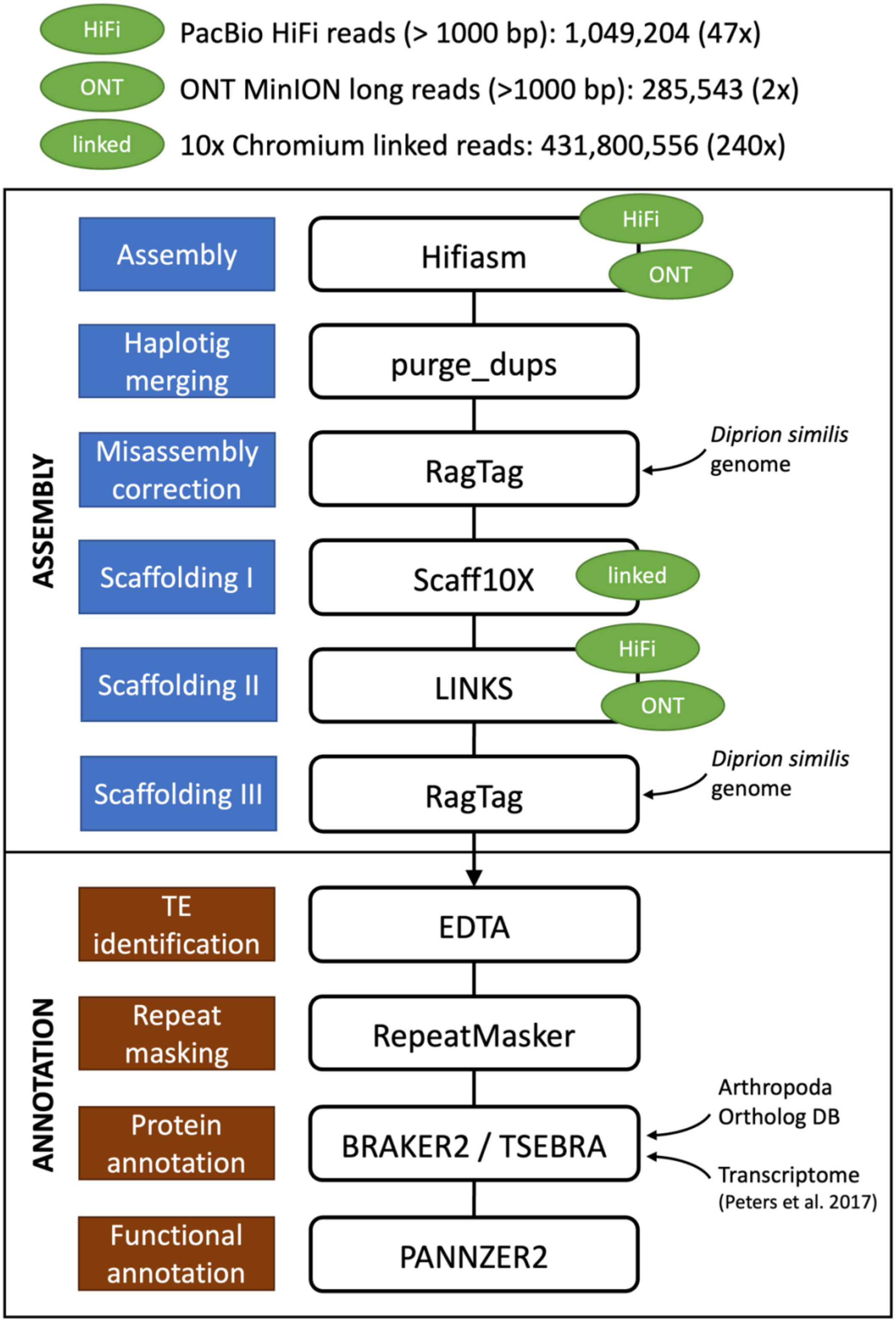


**Figure S1.** Workflow visualization of the genome assembly and annotation pipeline.

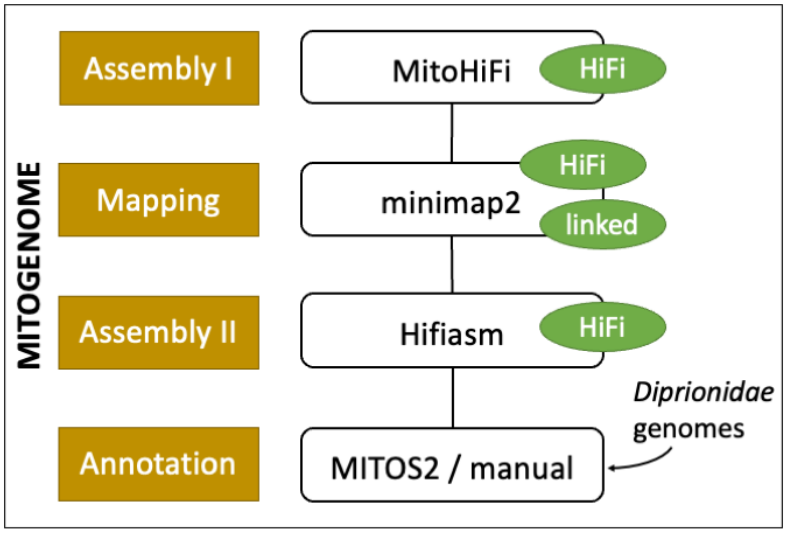


**Figure S2.** Workflow visualization assembling and annotating the mitochondrial genome.

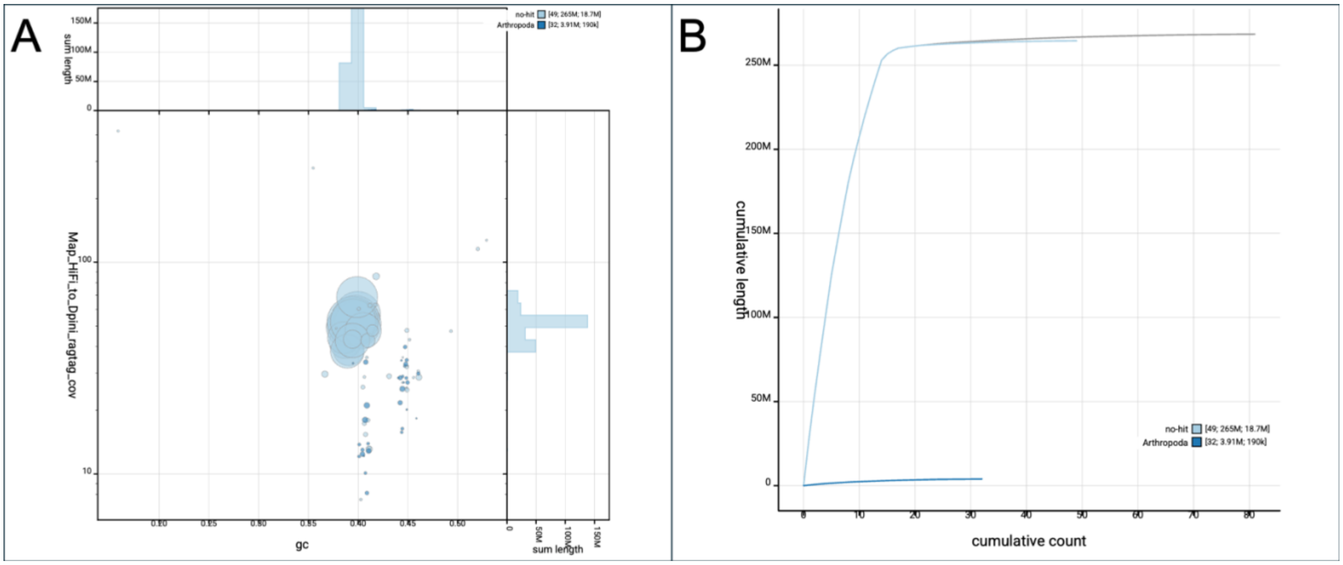


**Figure S3.** BlobToolKit assembly views. (A) Blob plot showing the distribution of assembly scaffolds on GC proportion and coverage; (B) Cumulative assembly span plot showing curves for subsets of scaffolds assigned to each phylum relative to the overall assembly.

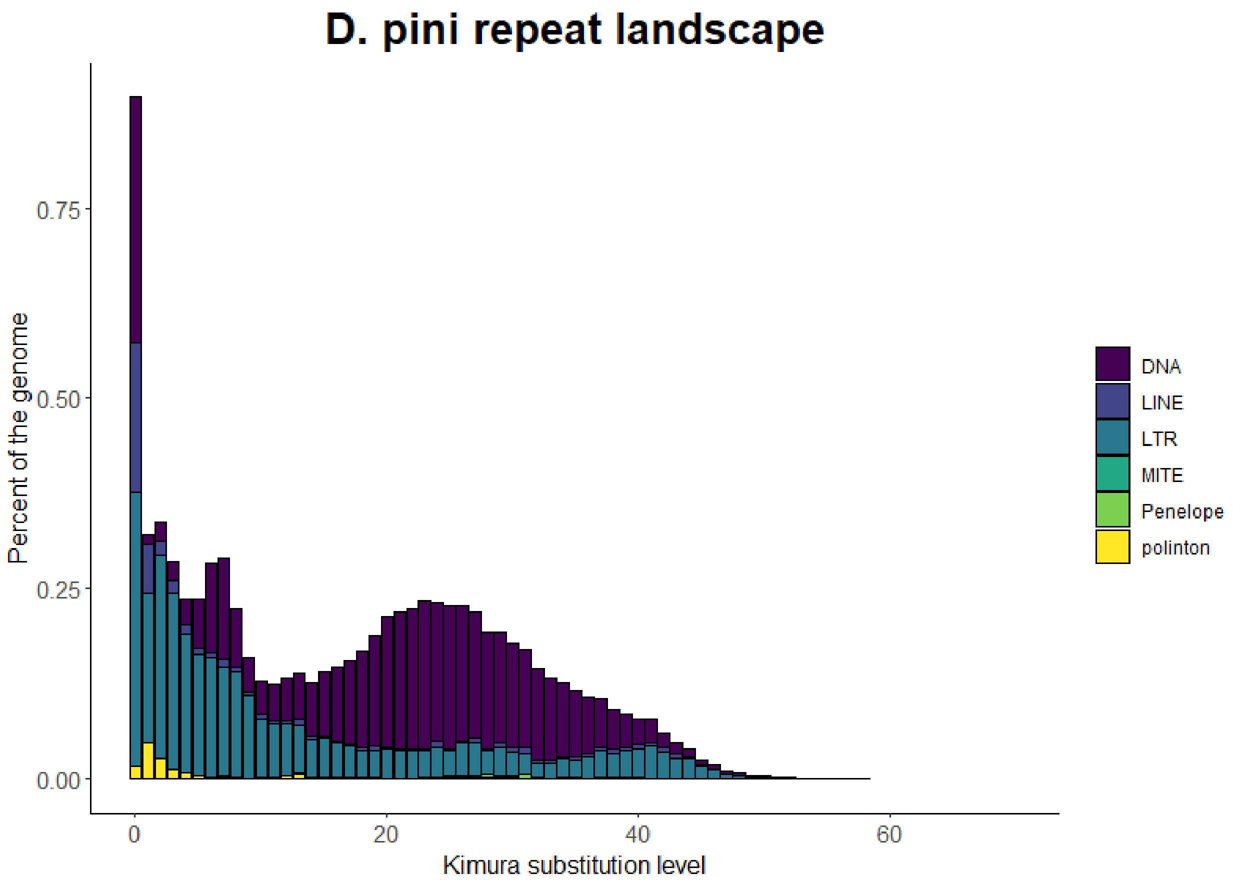


**Figure S4.** Repeat landscape.

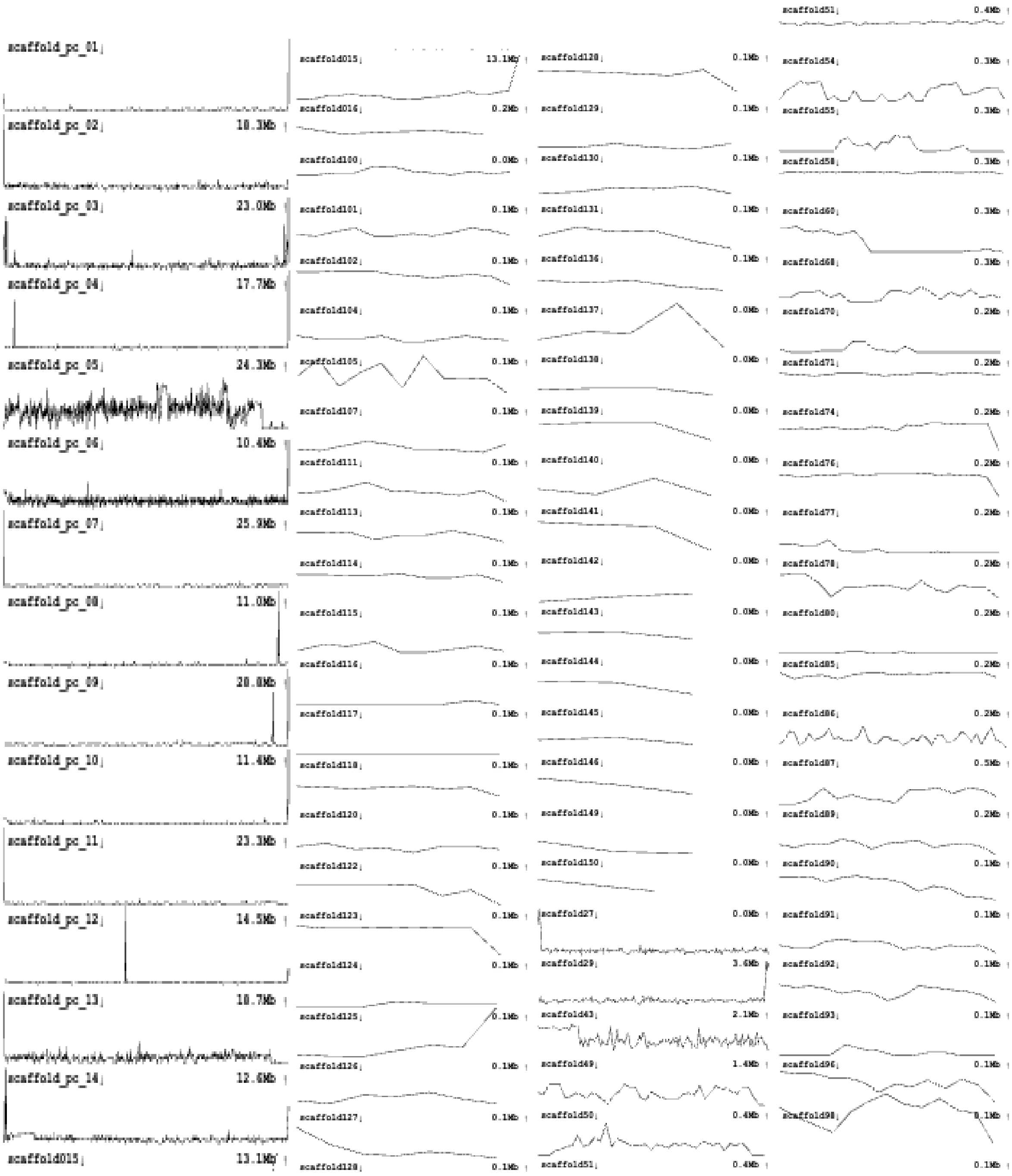

**Figure S5.** Occurrence of the common arthropod repeat $(TTAGG)_n$ within the assembled scaffolds.

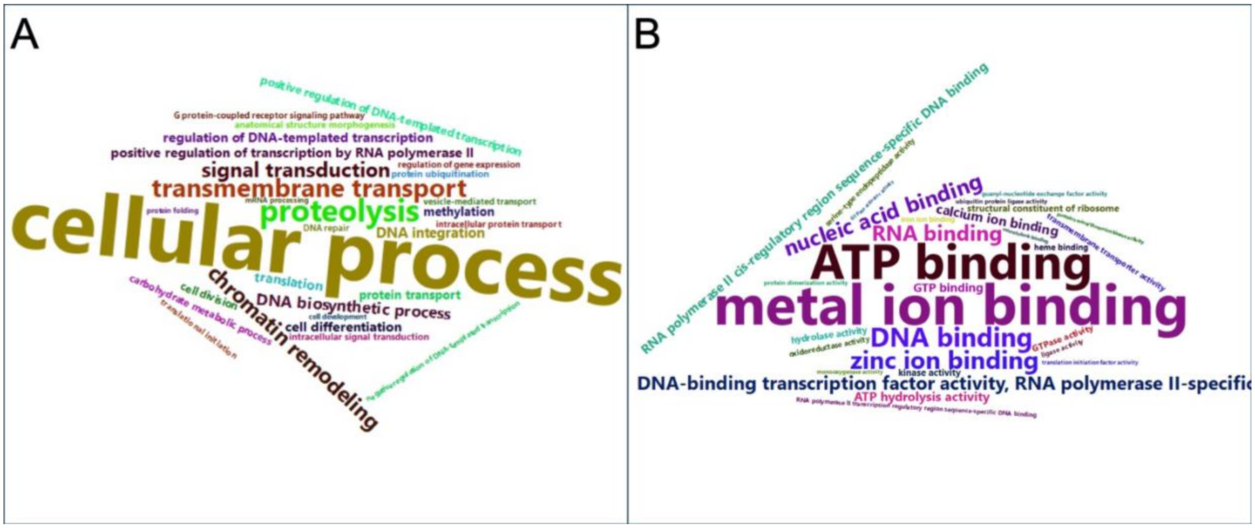


**Figure S6.** Word clouds of the 30 most abundant gene ontology (GO) terms in the *D. pini* complete protein set. (A) Annotations of biological processes (BP); (B) Annotations of molecular functions (MF).

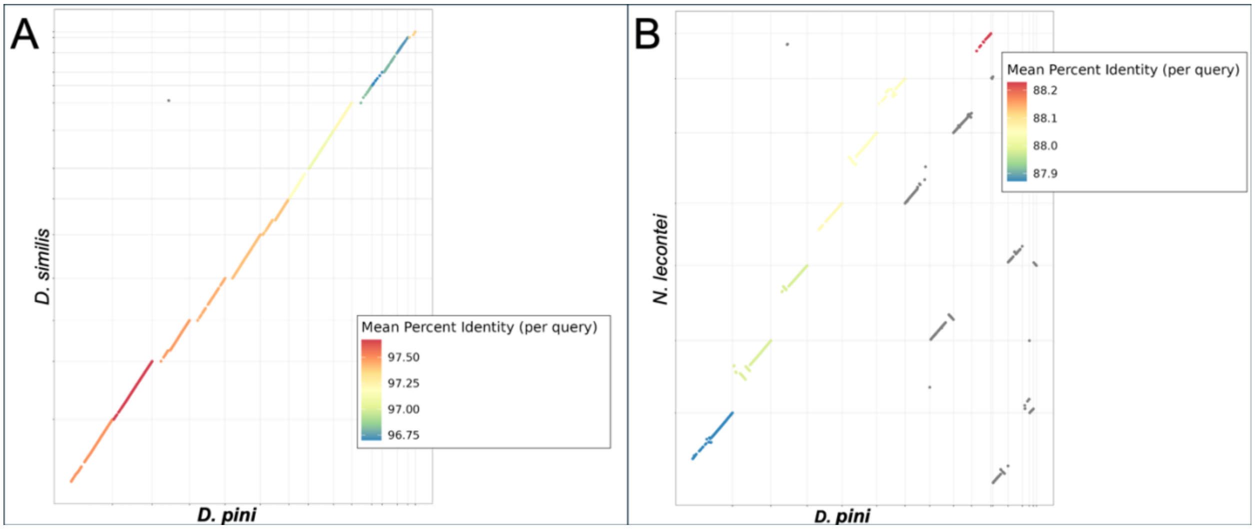


**Figure S7.** Dot plots of whole-genome alignments: (A) *D. pini* vs. *D. similis*, (B) *D. pini* vs. *N. lecontei*.
